# Variational autoencoders for cancer data integration: design principles and computational practice

**DOI:** 10.1101/719542

**Authors:** Nikola Simidjievski, Cristian Bodnar, Ifrah Tariq, Paul Scherer, Helena Andres-Terre, Zohreh Shams, Mateja Jamnik, Pietro Liò

## Abstract

International initiatives such as the Molecular Taxonomy of Breast Cancer International Consortium (METABRIC) are collecting multiple data sets at different genome-scales with the aim to identify novel cancer bio-markers and predict patient survival. To analyse such data, several machine learning, bioinformatics and statistical methods have been applied, among them neural networks such as autoencoders. Although these models provide a good statistical learning framework to analyse multi-omic and/or clinical data, there is a distinct lack of work on how to integrate diverse patient data and identify the optimal design best suited to the available data.

In this paper, we investigate several autoencoder architectures that integrate a variety of cancer patient data types (e.g., multi-omics and clinical data). We perform extensive analyses of these approaches and provide a clear methodological and computational framework for designing systems that enable clinicians to investigate cancer traits and translate the results into clinical applications. We demonstrate how these networks can be designed, built and, in particular, applied to tasks of integrative analyses of heterogeneous breast cancer data. The results show that these approaches yield relevant data representations that, in turn, lead to accurate and stable diagnosis.

## 1 Introduction

The rapid technological developments in cancer research yield large amounts of complex heterogeneous data on different scales – from molecular to clinical and radiological data. The limited number of samples that can be collected are usually noisy, incompletely annotated, sparse and high-dimensional (many variables). As much as these high-throughput data acquisition approaches challenge the data-to-discovery process, they drive the development of new sophisticated computational methods for data analysis and interpretation. In particular, the synergy of cancer research and machine learning has led to groundbreaking discoveries in diagnosis, prognosis and treatment planning for cancer patients (Levine et al., 2019; Vial et al., 2018). Typically, such machine learning methods are developed to address particular complexities inherent in individual data types, separately. While relevant, this approach is sub-optimal since it fails to exploit the inter-dependencies between the different data silos, and is thus often not extendable to analysing and modelling more complex biological phenomena (Hériché et al., 2019; Gomez-Cabrero et al., 2014).

To capitalise on the inter-dependencies and relations across heterogeneous types of data about each patient (Yuan et al., 2011; Miotto et al., 2016), integrating multiple types and sources of data is essential.The data-integration paradigm focuses on a fundamental concept – that a complex biological process is a combination of many simpler processes and its function is greater than the sum of its parts. Hence, integrating and simultaneously analysing different data types offers better understanding of the mechanisms of a biological process and its intrinsic structure. Many studies have addressed and highlighted the importance of data integration at different scales (Žitnik et al., 2019; López de Maturana et al., 2019; Karczewski and Snyder, 2018; Huang et al., 2017; Gomez-Cabrero et al., 2014). In the context of analysing cancer data, it has been shown that such integrative approaches yield improved performance for accurate diagnosis, survival analysis and treatment planning (Vial et al., 2018; Gevaert et al., 2006; Thomas et al., 2014; Kristensen et al., 2014; Shen et al., 2009). In particular, Wang et al. (2014) show that, for the case of 5 different cancer profiles, integrating mRNA expression, DNA methylation and miRNA data leads to more accurate survival profiles than each of the individual types of data alone. These findings are in line with the ones of (Amin et al., 2014), where the authors point out that gene expression profiles alone are sub-optimal for predicting complete response in patients with multiple myeloma.

In this paper we design and systematically analyse several deep-learning approaches for data integration based on Variational Autoencoders (VAEs) (Kingma and Welling, 2014). VAEs provide an *unsupervised* methodology for generating meaningful (disentangled) latent representations of integrated data. Such approaches can be utilised in two ways. First, the generated latent representations of integrated data can be exploited for analysis by any machine learning technique. Second, our architectures can be deployed on other heterogeneous data sets. We illustrate the functionality and benefit of the designed approaches by applying them to cancer data – this paves the way to improve survival analysis and bio-marker discovery.

There are several existing machine learning approaches that integrate diverse data. These can be classified into three different categories based on how the data is being utilised (Gevaert et al., 2006; Pavlidis et al., 2002): (i) output (or late) integration, (ii) partial (or intermediate) integration and (iii) full (or early) integration. Output integration relates to methods that model different data separately, the output of which is subsequently combined (Gevaert et al., 2006; Yang et al., 2010; Qi, 2012). Partial integration refers to specifically designed and developed methods that produce a joint model learned from multiple data simultaneously (Wang et al., 2014; Žitnik and Zupan, 2015; Gevaert et al., 2006). Finally, full-integration approaches focus on combining different data before applying a learning algorithm, either by simply aggregating them or learning a common latent representation (Shen et al., 2009; Bengio et al., 2013). Our work presented here falls into this third category, namely full (or early) integration.

Recently, many deep learning approaches have been proposed for analysing cancer data (Levine et al., 2019). Typically, they rely on extracting valuable features using deep convolutional neural networks for analysing and classifying tasks of radiological data (Esteva et al., 2019; Ardila et al., 2019). However, these methods often relate to supervised learning, and require many labelled observations in order to perform well. In contrast, unsupervised approaches learn representations by identifying patterns in the data and extracting meaningful knowledge while overcoming data complexities. Particular variants of deep learning networks, referred to as autoencoders, have demonstrated good performance for unsupervised representation learning (Bengio et al., 2013).

Autoencoders learn a compressed representation (embedding/code) of the input data by reconstructing it on the output of the network. The hope is that such a compressed representation captures the structure of the data (i.e., intrinsic relationships between the data variables) and therefore allows for more accurate downstream analyses (Belkin and Niyogi, 2003). Autoencoders have been deployed on a variety of tasks across different data types such as dimensionality reduction, data denoising, compression and data generation. In the context of cancer data integration, several studies highlighted their utility in combining data on different scales for identifying prognostic cancer traits such as liver (Chaudhary et al., 2018), breast (Tan et al., 2015) and neuroblastoma cancer (Zhang et al., 2018) sub-types. The focus of these studies is to apply autoencoders to specific problems of cancer-data integration.

In contrast, in this paper we investigate approaches that build upon probabilistic autoencoders which implement Variational Bayesian inference for unsupervised learning of latent data representations. Instead of only learning a compressed representation of the input data, VAEs learn the parameters of the underlying distribution of the input data. VAEs can be utilised as methods for full/early integration of data: this allows for learning representations from heterogeneous data on different scales from different sources. In this paper we mainly focus on the data integration aspect, so we utilise VAEs together with other sophisticated machine learning methods for modelling and analysing breast cancer data. We perform a systematic evaluation (we evaluate 1296 different network configurations) of different aspects of data integration based on VAEs. We investigate and evaluate four different integrative VAE architectures and their components. We analyse and demonstrate their functionality by integrating multi-omics and clinical data for different breast-cancer analysis tasks on data from the Molecular Taxonomy of Breast Cancer International Consortium (METABRIC) cohort. In summary, the contribution of this paper is two-fold: (i) novel architectures for integrating data; and (ii) methodologies for choosing architectures that best suit the data in hand.

## 2 Materials and Methods

Many machine learning methodologies have been applied to cancer medicine to improve and personalise diagnosis, survival analysis and treatment of cancer patients. These include linear and non-linear, as well as supervised and unsupervised techniques like regression, PCA, SVMs, deep neural nets, and autoencoders (Kourou et al., 2015).

Some are more suitable for integrating diverse types of data than others. In our work we use VAEs and combine them into a number of different architectures for a deep analysis and comparison with respect to specific data features and tasks at hand. VAEs are particularly suitable in this setting since they are generative, non-linear, unsupervised and amenable to integrating diverse data.

We deploy our architectures on the case of integrating multi-omic and clinical cancer data. There are a number of candidate initiatives for big data collection of cancer data such as TCGA and METABRIC. In our work we use the METABRIC data set because it is one of the largest amongst genetic data sets, it is reasonably well annotated, and it is well analysed. We particularly focus on the integration of gene expressions, copy number alterations and clinical data.

In this section we describe theoretical aspects of VAEs and the specialised architectures that we use to integrate data. Next, we describe the data and the suite of experiments used to evaluate the methodological and computational frameworks for investigating cancer traits in clinical applications.

### 2.1 Variational Autoencoders

Generally, an autoencoder consists of two networks, an *encoder* and a *decoder*, which broadly perform the following tasks:

- **Encoder**: Maps the high dimensional input data into a latent variable embedding which has lower dimensions than the input.
- **Decoder**: Attempts to reconstruct the input data from the embedding.

The model contains a decoder function *f*(·) parameterised by *θ* and an encoder function *g*(·) parameterised by *ϕ*. The lower dimensional embedding learned for an input *x* in the bottleneck layer is *h* = *g*_*ϕ*_(*x*) and the reconstructed input is *x*′ = *f*_*θ*_(*g*_*ϕ*_ (*x*)).

The parameters ⟨*θ, ϕ*⟩ are learned together to output a reconstructed data sample that is ideally the same as the original input *x* ≈ *f*_*θ*_(*g*_*ϕ*_ (*x*)). There are various metrics used to quantify the error between the input and output such as cross entropy (CE) or simpler metrics such as mean squared error:

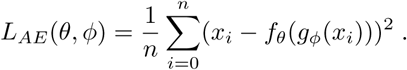

The main challenge when designing an autoencoder is its sensitivity to the input data. While an autoencoder should learn a representation that embeds the key data traits as accurately as possible, it should also be able to encode traits which generalise beyond the original training set and capture similar characteristics in other data sets.

Thus, several variants have been proposed since autoencoders were first introduced. These variants mainly aim to address shortcomings such as improved generalisation, disentanglement and modification to sequence input models. Some significant examples include the Denoising Autoencoder (DAE) (Vincent et al., 2008), Sparse Autoencoder (SAE) (Coates et al., 2011; Makhzani and Frey, 2014), and more recently the Variational Autoencoder (VAE) (Kingma and Welling, 2014).

The VAE (Figure 1) uses stochastic inference to approximate the latent variables *z* as probability distributions. These distributions represent and capture relevant features from the input. VAEs are scalable to large data sets, and can deal with intractable posterior distributions by fitting an approximate inference or recognition model, using a reparameterised variational lower bound estimator. They have been broadly tested and used for data compression or dimensionality reduction. Their adaptability and potential to handle non-linear behaviour has made them particularly well suited to work with complex data.

**Figure 1:**
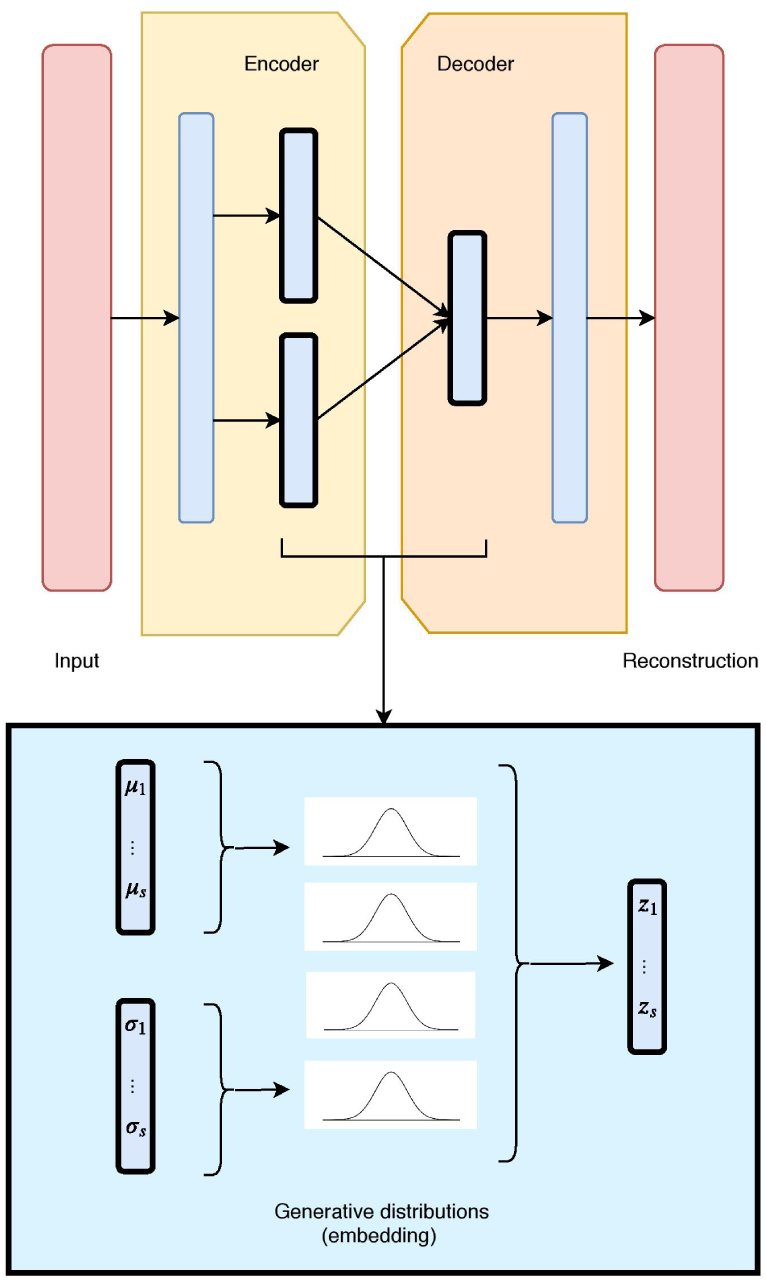
The unimodal VAE architecture and latent embedding: the red layers correspond to the input and reconstructed data, given and generated by the model. The hidden layers are in blue, with the embedding framed in black. Each latent component is made of two nodes (mean and standard deviation), which define a Gaussian distribution. The combination of all Gaussian constitutes the VAE generative embedding.

A VAE builds upon a probabilistic framework where the high dimensional data *x*is drawn from a random variable with distribution *p*_*data*_(*x*). It assumes that the natural data *x* also lies in a lower dimensional space, that can be characterised by an unobserved continuous random variable *z*. In the Bayesian approach, the prior *p*_*θ*_(*z*) and conditional (or likelihood) *p*_*θ*_(*x*|*z*) typically come from a family of parametric distributions, with Probability Density Functions (PDFs) differentiable almost everywhere with respect to both *θ* and *z*. While the true parameters *θ* and the values of the latent variables *z* are unknown, the VAE approximates the often intractable true posterior *p*_*θ*_(*z*|*x*) by using a recognition model *q*_*ϕ*_ (*z*|*x*) and the learned parameters *ϕ* represented by the weights of a neural network.

More specifically, a VAE builds an inference or a recognition model *q*_*ϕ*_ (*z*|*x*), where given a data-point *x* it produces a distribution over the latent values *z* from where it could have been drawn. This is also called a probabilistic encoder. A probabilistic decoder will then, given a certain value of *z*, produce a distribution over the possible corresponding values of *x*, therefore constructing the likelihood *p*_*θ*_(*x*|*z*). Note that the decoder is also a generative model, since the likelihood *p*_*θ*_(*x*|*z*) can be used to map from the latent to the original space and learn to reconstruct the inputs as well as generate new ones.

Typically, VAE model assumes latent variables to be the centred isotropic multivariate Gaussian *p*_*θ*_(*z*) = *N* (*z*; 0, *I*), and *p*_*θ*_(*x*|*z*) a multivariate Gaussian (for numerical values) or Bernoulli (for categorical values) with parameters approximated by using a fully connected neural network. Since the true posterior *p*_*θ*_(*z*|*x*) is intractable, we assume it takes the form of a Gaussian with an approximately diagonal covariance. This allows the variational inference alternative to approximate the true posterior, as it converts the inference problem into an optimisation one. In particular, instead of solving intractable integrals, this relates to maximising a likelihood. In such cases, the variational approximate posterior will also need to be a multivariate Gaussian with diagonal covariate structure:

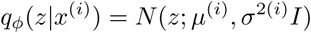

where the mean *µ*^(*i*)^ and standard deviation *σ*^(*i*)^ are outputs of the encoder.

Since *p*_*θ*_(*z*) and *q*_*ϕ*_ (*z*|*x*^(*i*)^) are Gaussian, the discrepancy between them can be directly computed and differentiated. The resulting likelihood for this model on data-point *x*^(*i*)^ is:

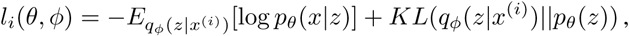

where the first term corresponds to the reconstruction loss, which encourages the decoder to learn to reconstruct the data from the embedding space. The second term is regularisation, and measures the divergence between the encoding distributions *q*(*z*|*x*) and *p*(*z*), and penalises the entanglement between components in the latent space. It is typically estimated by the Kullback-Leibler (KL) divergence, a measure of discrepancy between two probability distributions, which in this case is applied between the prior and the representation.

While in this paper we focus on a standard Gaussian prior due to its simplicity, there are a severeal, more sophisticated, alternatives for the choice of a prior. In particular, Dilokthanakul et al. (2016) propose a mixture of Gaussians in order to achieve more flexible priors, and Tomczak and Welling (2018) realise this by estimating the prior as a mixture of approximate posteriors. Nalisnick and Smyth (2016) employ a Dirichlet process as a non-parametric prior through stick-breaking process, which generalises over the generative process and allows for better representations. Johnson et al. (2016) utilise graphical models as a prior to train a VAE model. These alternative approaches to the choice of a prior require more sophisticated model training techniques in the learning phase. On the other hand, there are also approaches that instead of the prior, they focus on more flexible posteriors, therefore leading to better (and disentangled) representations. These include normalising flows (Rezende and Mohamed, 2015), auto-regressive flows (Chen et al., 2017) and inverse auto-regressive flows (Kingma et al., 2016).

In a similar context, research has shown that the entanglement factor can play a crucial role in the quality of the representations. In response, Higgins et al. (2017) control the influence of the disentanglement factor using a parameter *β*. Moreover, some approaches have experimented with different regularisation terms, such as the InfoVAE (Zhao et al., 2017), where Maximum Mean Discrepancy (MMD) is employed as an alternative to KL divergence. MMD (Gretton et al., 2007) is based on the concept that two distributions are identical if, and only if, all their moments are identical. Therefore, by employing MMD via the kernel embedding trick, the divergence can be defined as the discrepancy between the moments of two distributions *p*(*z*) and *q*(*z*) as:

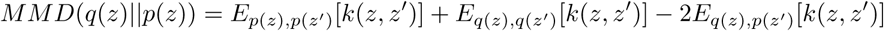

where *k*(*z, z*′) denotes any universal kernel. In this paper, we employ a Gaussian kernel 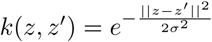, when considering MMD regularisation in the objective function.

### 2.2 Variational Autoencoders for data integration

We designed and evaluated four different architectures for data integration: we present them here each with two diverse data sources (depicted in Figures 2, 3, 4 and 5 as red and green boxes on the left).

**Figure 2:**
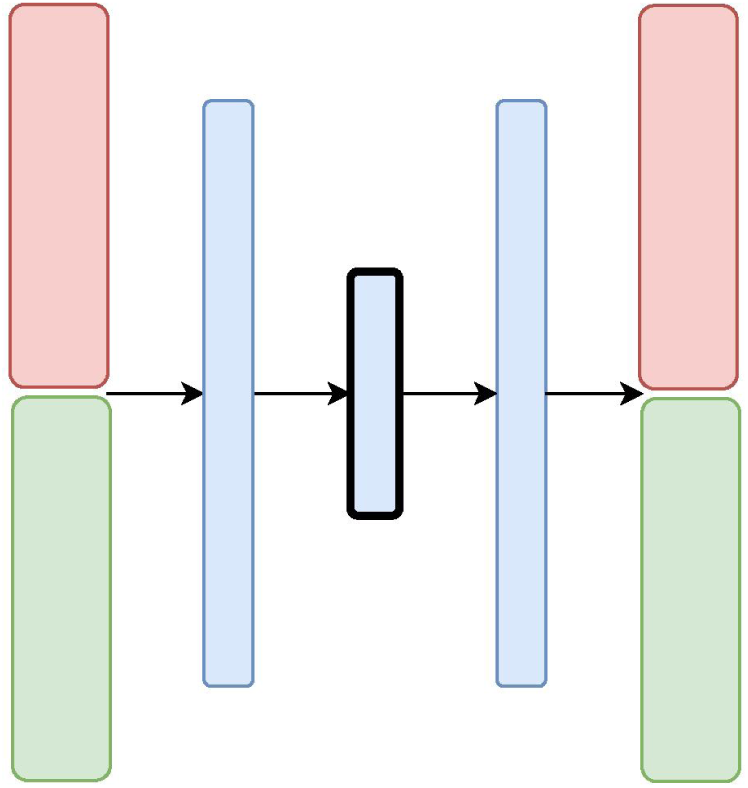
The CNC-VAE Architecture: the red and green layers on the left correspond to two inputs from different data sources. The blue layers are shared, with the embedding being framed in black.

**Figure 3:**
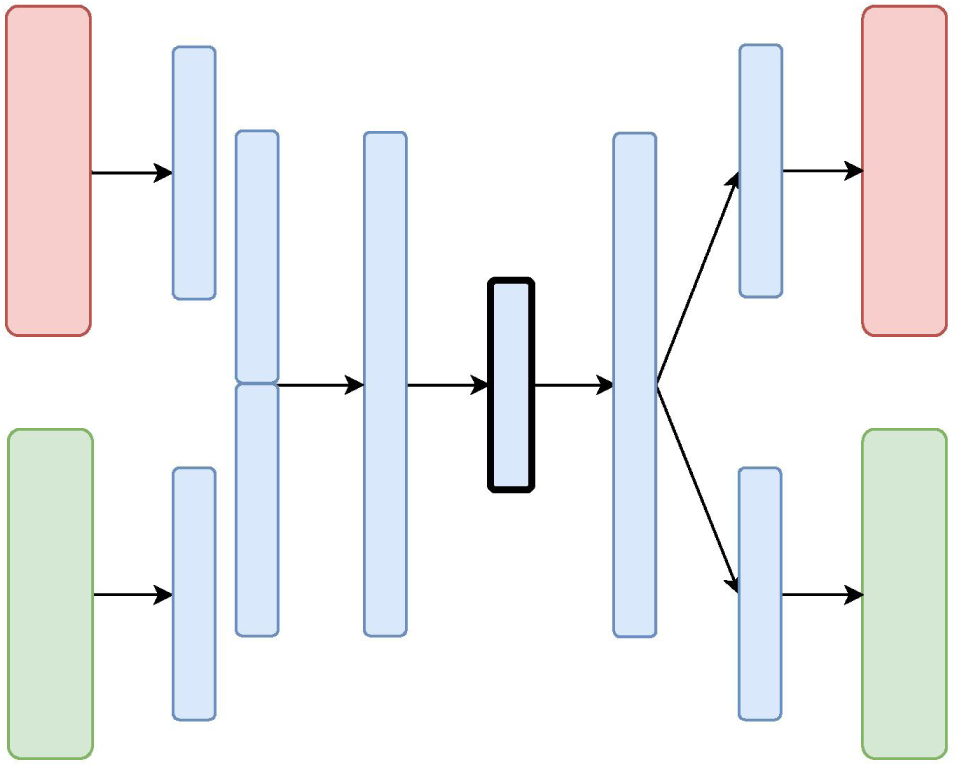
The X-VAE Architecture: the red and green layers on the left correspond to two inputs from different data sources. The blue layers are shared, with the embedding being framed in black.

**Figure 4:**
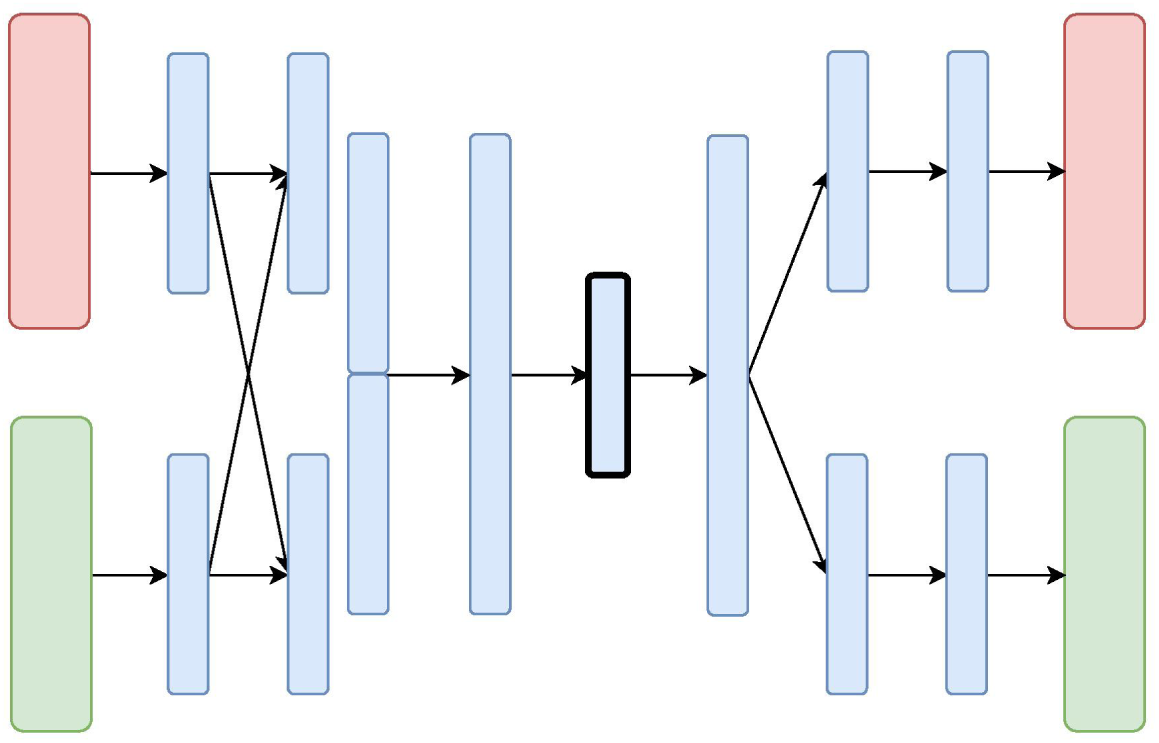
The MM-VAE Architecture: the red and green layers on the left correspond to two inputs from different data sources. The blue layers are shared, with the embedding being framed in black.

**Figure 5:**
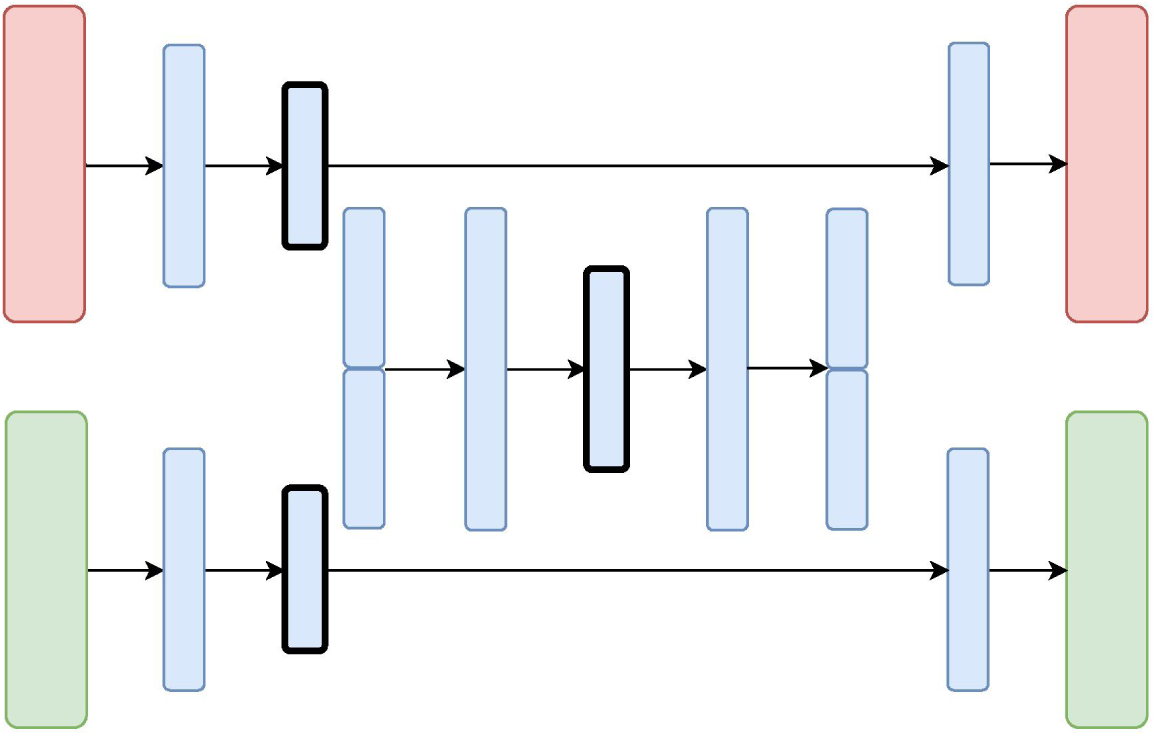
The H-VAE Architecture: the red and green layers on the left correspond to two inputs from different data sources. The blue layers are shared, with the embedding being framed in black.

The first architecture, **Variational Autoencoder with Concatenated Inputs (CNC-VAE)** in Figure 2, is a simple approach to integration, where the encoder is directly trained from different data sets, aligned and concatenated at input. While such architecture is a straightforward and not a novel way to data integration, we employ it both, as a benchmark and a proof-of-principle for learning a homogeneous representation from heterogeneous data sources.

Besides the concatenated input, the rest of the CNC-VAE network utilises a standard VAE architecture. As depicted in Figure 2, the input data is first scaled, aligned and concatenated before being fed to the network. CNC-VAE has one objective function that reconstructs the combined data rather than a separate objective function for each input data source. Therefore, CNC-VAE aims at reducing redundancies and extracting meaningful structure across all input sources, regardless of the scales or modalities of the data. Whilst the CNC-VAE architecture may be simplistic, the complexity lies in highly domain-specific preprocessing of the data. Indeed, in some real-world settings, utilising a single objective function of combined heterogenious inputs may not be optimal or even feasible.

Unlike CNC-VAE, the next three architectures aim at more sophisticated means to data integration. In particular, all of them consider data integration in the hidden layers. The **X-shaped Variational Autoencoder (X-VAE)** merges high-level representations of several heterogeneous data sources into a single latent representation by learning to reconstruct the input data from the common homogeneous representation. The architecture is depicted in Figure 3 and consists of individual branches (one for each data source: red and green) that are combined into one before the bottleneck layer. In the decoding phase, the merged branch splits again into several branches that produce individual reconstructions of the inputs. X-VAE takes into account different data modalities by combining different loss functions for each data source in the objective function. This allows for learning better and more meaningful representations.

While, in principle, X-VAE is able to take into account many possible interactions between multiple data sources, its performance is sensitive to the properties of the data being integrated. In particular, X-VAE is prone to poor performance when employed to integrate unbalanced data sets with low number of observations. As a consequence, the objective function might also be unbalanced, focusing on some sources more if the distribution of the input data varies substantially across the data sources. A similar limitation can also result from a poor choice of loss function for each of the data sources.

The **Mixed-Modal Variational Autoencoder (MM-VAE)** attempts to address some of the limitations of X-VAE, by employing a more gradual integration in the hidden layers of the encoder. More specifically, it builds upon the concept of transfer learning, where learned concepts from one domain are re-purposed and shared for learning tasks in others domains. Figure 4 presents the architecture of MM-VAE. Similarly to X-VAE, it also consists of branches that individually reconstruct the input data sources. Here, however, the important difference is that the branches share information with each other in the encoding phase. In particular, higher-level learned concepts of each branch are shared between all the branches, and used deeper in the network. This allows for information from the different sources to be combined more gradually before being compressed into a single homogeneous embedding.

The objective function combines different reconstruction loss functions that correspond to the data types at input. Similarly to X-VAE, MM-VAE’s performance is limited when small and unbalanced data sets are being considered. While the additional integration layers may help to stabilise the objective function, poor choice of reconstruction loss terms may still impede the performance in general.

The **Hierarchical Variational Autoencoder (H-VAE)** builds upon traditional meta-learning approaches for combining multiple individual models. H-VAE, depicted in Figure 5, is comprised of several low-level VAEs that relate to each data source separately, and the result is assembled together in a high-level VAE. More specifically, each of the low-level VAEs is employed to learn a representation of an individual data source. These individual representations are then merged together and fed to a high-level VAE that produces the integrated data representation. We use the same architecture for each low-level VAE, but in principle, these could be independently designed and further refined for a specific data-source and task at hand.

H-VAE is designed to improve on some of the shortcomings of X-VAE and MM-VAE, since it simplifies the individual network branches. In particular, the input to the high-level autoencoder is composed of representations learned from several individual low-level autoencoders. These low-level autoencoders already implement distribution regularisation terms in each of them separately, thus the input to the high-level autoencoder already consists of approximated multivariate standard normal distributions characterising the general traits of the individual input modalities. Moreover, since each data source is handled in a modular fashion, H-VAEs are capable of handling data sets which make best use of specialised low-level autoencoders. However, constructing an H-VAE adds a substantial computational overhead compared to the other three architectures as it involves a two-stage learning process where low-level VAEs must be trained first, and then the final high-level representation can be learned on the outputs of the low-level encoders.

### 2.3 Data

To demonstrate how the proposed VAE architectures can be utilised in the integration of heterogeneous cancer data types, we conducted our study utilising multi-omics data found on somatic copy number aberrations (CNA), mRNA expression data, as well as on the clinical data of breast cancer patient samples from the Molecular Taxonomy of Breast Cancer International Consortium (METABRIC) cohort (Curtis et al., 2012).

Providing effective treatment takes such heterogeneity of data into account, and our VAE architectures enable us to do just that. Finding driver events which help stratify breast cancers into different subgroups has been of great focus within the research community lately, particularly the identification of genomic profiles that stratify patients.

In the context of genomic and transcriptomic studies, the acquired somatic mutations and the inherited genomic variation contribute jointly to tumorigenesis, disease onset and progression (Curtis et al., 2012; Tan et al., 2015; Pereira et al., 2016). For example, despite somatic copy number aberrations being the dominant feature found in sporadic breast cancer cases, the elucidation of driver events in tumorigenesis is hampered by the large variety of random non-pathogenic passenger alterations and copy number variants (Leary et al., 2008; Bignell et al., 2010).

This has led to the argument that integrative approaches for the available information are necessary to make richer assessments of disease sub-categorisation (Curtis et al., 2012). A pioneering work that advocates this perspective in breast cancer research is the METABRIC initiative. The METABRIC project is a Canada-UK initiative that aims to group breast cancers based on multiple genomic, transcriptomic and image data types recorded over 2000+ patient samples. This data set represents one of the largest global studies of breast cancer tissues performed to date. Similarly to (Curtis et al., 2012) we focus on integrating CNA and mRNA expression data, but in addition integrate clinical data too. We use integrative VAEs to showcase how such architectures can be designed, built and used for cancer studies of this kind.

### 2.4 Experimental setup

What follows is an outline of our experimental evaluation used to verify that the studied approaches produce valid representations and can be employed for data integration. The aim of this evaluation is threefold. First, for each of the architectures, we seek the optimal configuration in terms of choosing an appropriate objective function and parameters of the network. Second, we aim to evaluate and choose the most appropriate architectures for our data-integration tasks. In particular, we perform a comparative quantitative analysis of the representations obtained from each of the architectures based on different data sets at input. Finally, we discuss the findings in terms of their application to cancer data integration and provide a qualitative (visual) analysis of the obtained representations.

In particular, we tackle several classification tasks by integrating three data types from the METABRIC data - Copy Number Alterations (CNA), mRNA expression and clinical data. We evaluate the predictive performance of the integrative approaches by combining clinical and mRNA data, CNA and mRNA data as well as clinical and CNA data, separately. The METABRIC data consists of 1,980 breast-cancer patients assigned to different groups according to:

- two immunohistochemistry (IHC) sub-types (ER+ and ER-),
- six intrinsic gene-expression sub-types (PAM50) (Prat et al., 2010), and
- ten IntegrativeCluster (IntClust) sub-types (Curtis et al., 2012).

These patients are also assigned to two groups based on whether or not the cancer metastasised to another organ after the initial treatment (i.e., Distance Relapse). The three cancer sub-types and the distance relapse variable (described with gene expression profiles, CNA profiles and clinical variables for each patient), are used as target variables in the classification tasks performed in the study.

To control our study, we followed Curtis et al. (2012) and used a pre-selected set of the input CNA and mRNA features. In particular, we used the most significant *cis*-acting genes that are significantly associated with CNAs determined by a gene-centric ANOVA test. We selected the genes with the most significant Bonferroni adjusted p-value from the Illumina database containing 30566 probes. After missing-data removal, the input data sets consisted of 1000 features of normalised gene expression numerical data, scaled to [0, 1], and 1000 features of copy number categorical data. The clinical data included various categorical and numerical features such as: age of the patient at diagnosis, breast tumour laterality, the Nottingham Prognostic Index, inferred menopausal state, number of positive lymph nodes, size and grade of the tumour as well as chemo-, hormone- and radio-therapy regimes. Numerical features were discretised and subsequently one-hot encoded. This was combined with the categorical features, yielding 350 clinical features. Finally, all three data sets were sampled into 5-fold cross-validation splits for each classification tasks separately, stratified according to the class distribution of the 4 target variables, respectively. Note that these splits remained the same for all experiments in the study.

While our four architectures differ in some key aspects related to how and where (on which level) they integrate data, for experimental purposes of this study, the depth of the architectures remained moderate and constant across all experiments. In particular, in all designs except for MM-VAE, the encoder and decoder were symmetric and consisted of compression/decompression dense layers placed before and after data merging. MM-VAE implemented an additional data-merging layer in the encoder network. Therefore, all of the architectures had a moderate depth between 2 and 4 hidden layers. The optimal output size of these layers was evaluated for different values of 128, 256 and 512. Moreover, all layers used batch normalisation (Ioffe and Szegedy, 2015) with Exponential Linear Unit (ELU) (Clevert et al., 2015) activations (except for the bottleneck and the output layers). All of the architectures also employed a hidden dropout component with a rate of 0.2. Note that the final layers of the CNA and clinical branches employed sigmoid activation function. The models were trained for 150 epochs using an Adam optimiser (Kingma and Ba, 2014) with a learning rate of 0.001 (with exponential decay rates of first- and second-moment estimates *β*_1_ = 0.9 and *β*_2_ = 0.999) and a batch size of 64. Furthermore, we also investigated the performance of representations with different sizes. For each of the architectures and their configurations, we learned and evaluated representations with sizes 16, 32 and 64.

In the experiments we also considered choosing an optimal objective function that would improve the disentanglement between the embedded components. The objective functions consider both the reconstruction loss and a regularisation term. For the former, given that we integrated heterogeneous data, we incorporated Binary Cross Entropy (BCE) loss for the categorical and Mean Squared Error (MSE) loss for the continuous data. Note that, while the CNA data is categorical and so multivariate categorical distribution would be suitable, an approach such as one-hot encoding would substantially increase the data dimensionality. Therefore, we employed label smoothing (Salimans et al., 2016), where the form of *p*_***θ***_(***x***_*cna*_|*z*) is a multivariate Bernoulli distribution, with values of ***x***_*cna*_ scaled to [0, 1]. For the regularisation terms, we evaluated different options which include weighted KL divergence and weighted Maximum Mean Discrepancy (MMD). We tested different values of weight *β, β* ∈ {1, 10, 15, 25, 50, 100}, for each of the two regularisation terms.

To make optimal design decisions, we evaluated the quality of the representations obtained from our four integrative architectures on three integrative tasks, each of these with 108 different network configurations with respect to the hyper-parameters outlined above. In particular, we evaluated the performance of a given configuration by training a predictive model on the produced representations and measuring its predictive performance on a binary classification task of IHC cancer sub-types (ER+ and ER-). For all network configurations, we trained and evaluated a Gaussian naive Bayes classifier, since it does not require tuning of additional hyper-parameters for the downstream task. We performed a 5-fold cross-validation and report the average accuracy.

Once we identified the appropriate configuration for each of the architectures, we evaluated the quality of the learned representation in terms of predictive performance on the remaining three classification tasks. In particular, we evaluated the performance of three different methods trained on different representation. These included Gaussian naive Bayes classifier, SVMs (with RBF kernel *C* = 1.5 and gamma set to 1*/N*_*f*_, where *N*_*f*_ denotes the number of features) and Random Forest (with 50 trees and 1*/*2 of the features considered at every split). For all three classification tasks we also performed a 5-fold cross-validation and report the average accuracy. We also compared these results with the performance of predictive models trained on: (i) the raw (un-compressed) data, as well as (ii) data transformed using PCA (a linear method for data transformation).

The integrative VAE architectures are implemented using the Keras deep learning library (Chollet et al., 2015) with Tensorflow backend. The code for training and evaluating the performance of the VAE networks is available on this repository.^2^

Finally, we visually inspected the learned representations of the whole data set obtained from each of the architectures, and compared them to the uncompressed data. For this task we employed the t-distributed stochastic neighbouring embedding (tSNE) (van der Maaten and Hinton, 2008) algorithm.

## 3 Results

We present and discuss the results of the empirical evaluation. First, we report on the analyses for identifying the suitable design choices within the integrative approaches. Next, we present the results of the analyses of predictive performance of three different predictive methods applied to representations obtained from our VAE architectures with the optimal configuration. Finally, we present a visual analysis of the learned representations obtained from the evaluated architectures.

### 3.1 Design of integrative VAEs

For each integrative task, we investigated 108 different configurations for each architecture. These highlighted the effect of the size of the learned embedding, the optimal size of each of the dense layers, the most appropriate regularisation in the objective function, and how much this regularisation should influence the overall loss. We evaluated these configurations for all four architectures on three integrative tasks, by comparing the average train and test performance of classifying IHC sub-typed patients. The results, in general, indicate that properties of these configurations for each architecture are consistent across the three integrative tasks. Therefore, for brevity, here we only present the results when combining clinical and mRNA data. The rest of the results, namely for combining CNA and mRNA, and CNA and clinical data are given in the Supplementary Material.

Figure 6 presents the downstream performance of predictive models, trained on the representations produced by the integrative VAEs on clinical and mRNA data. In particular, Figure 6(a)-(d) compare the performance from representations obtained from CNC-VAE, X-VAE, MM-VAE and H-VAE, respectively. In general, the configurations regularised with MMD yield better representations that lead to substantially more accurate predictions than the configurations regularised with KL. In terms of the weight of the regularisation term, the configurations are robust in general, with moderately large weights (*β* = [25, 50]) leading to slightly better results.

**Figure 6:**
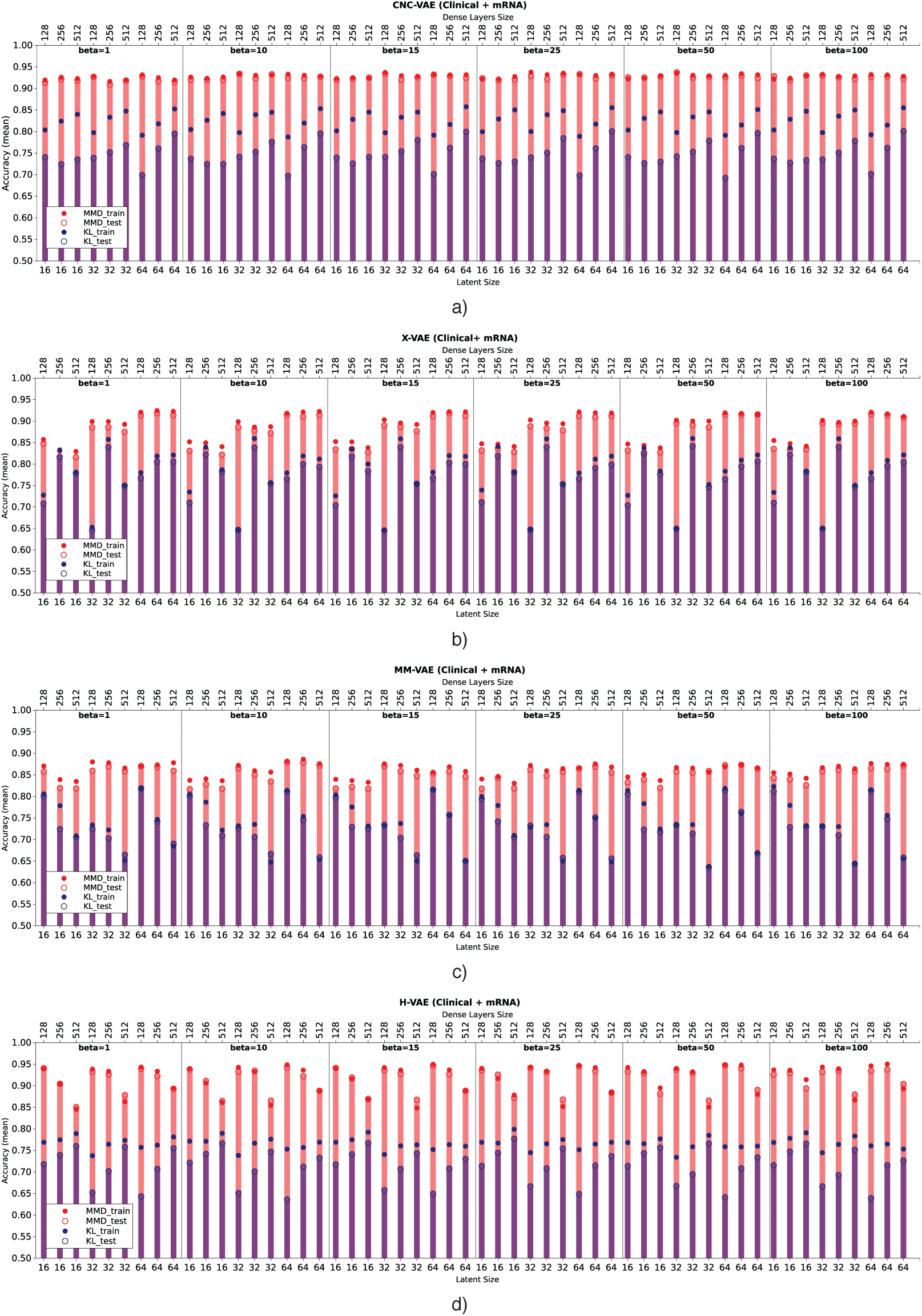
Comparison of the downstream performance on the IHC classification tasks of a predictive model trained on the representations produced by integrating clinical and mRNA data using (a) CNC-VAE (b) X-VAE (c) MM-VAE and (d) H-VAE. Full circles denote the training accuracy, while empty circles and bars denote the test accuracy averaged over 5-fold cross-validation. Red and blue colours denote the configurations when MMD and KL are employed, respectively. Bottom x-axis depicts the size of the latent dimension, while the top x-axis the size of the dense layers of each configuration.

In term of the size of the dense layers, all architectures except H-VAE exhibit stable behaviour, with moderate sizes of (*size* = [128, 256]) leading to slightly better representations than the ones with dense layer size of 512 in the case of X-VAE and MM-VAE. In the case of H-VAE, the quality of the representations is more affected by the size of the layer where smaller sizes lead to better performance than larger ones.

Considering the size of the latent space, the networks that produce higher-dimensional encodings lead to better predictive performance. This is particularly the case for X-VAE and MM-VAE architectures, while the other two are mostly unaffected. Note however, that the influence of the size of the representations on the overall performance is also related to the integrative task. More specifically, for this particular classification task, higher-dimensional representations when integrating clinical and mRNA data yield better and more stable performance overall. In contrast, when integrating clinical/CNA or CNA/mRNA data lower-dimensional representations are better.

In summary, based on these results, we made the following design decisions for configuring the integrative VAE architectures for the rest of the experimental analyses. First, the networks were trained using the MMD regularisation with *β* = 50, since in all cases using MMD exhibited better performance than the networks trained using KL divergence with various levels of *β*. Next, we set the size of the dense layers to 256. Finally, since large sizes of the latent space yielded better performance, we set it to be 64.

### 3.2 Quality of the learned representations

In this set of experiments, we focused on testing our central hypothesis that the integrative VAE architectures are able to produce representations that yield stable and improved predictive performance. We evaluated their performance in three classification tasks: predicting IC10, PAM50 sub-types and Distance Relapse.

We used three standard predictive methods: Naive Bayes, SVM and Random Forest. These were deployed: (i) on representations learned (compressed) from data integrated through our four VAE architectures; (ii) on embedded combined data using PCA with 64 components; (iii) on combined raw (un-compressed) data; and (iv) on each of the data sources separately in order to evaluate the integrative effect. Apart from this last case, the data sources for integration were CNA/mRNA, clinical/mRNA and clinical/CNA data, as before.

Table 1 summarises the results of this analysis. In general, all of the VAE integrative architectures outperform the baselines on all three predictive tasks when integrating CNA/mRNA, clinical/mRNA data and clinical/CNA. Overall, all architectures produce better representations when integrating clinical and mRNA data. This result is consistent across all three tasks, where the learned representations coupled with SVMs yield the best predictive performance. This finding is also supported by the benchmark approaches, where combining clinical and mRNA data yields better results than CNA/mRNA and clinical/CNA. Note that, for the task of predicting Distance Relapse, integrating clinical/CNA exhibits, in general, slightly worse but comparable performance to the one produced for clinical/mRNA. These results suggest that for our particular classification tasks, some data types are more beneficial to integrate than others.

**Table 1:**
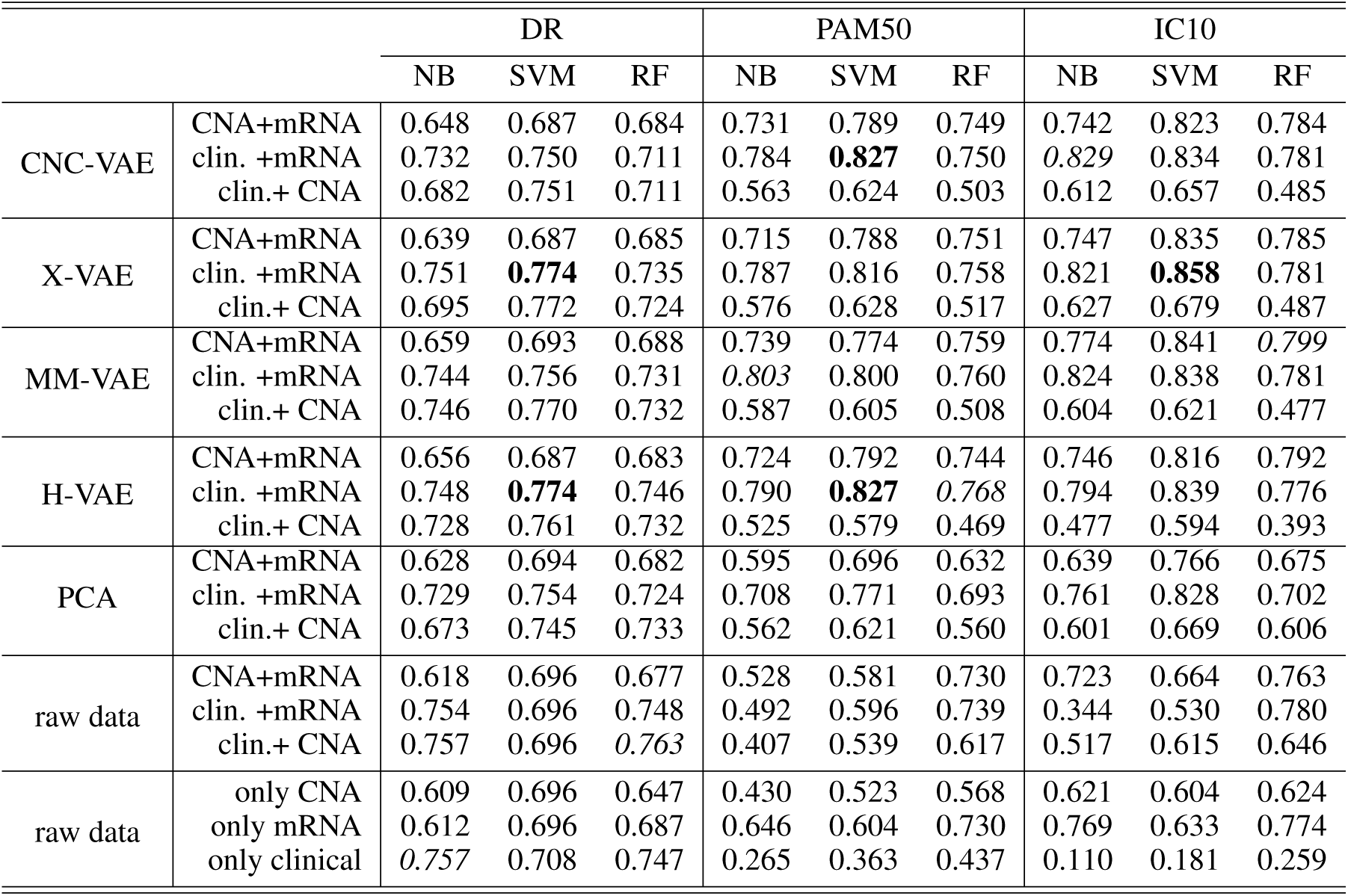
Comparison of the downstream predictive performance (on three classification tasks) of the three predictive models trained on raw and PCA-transformed data as well as representations produced by the four integrative VAEs by integrating CNA/mRNA, clinical/mRNA and clinical/CNA data. Italic typeface denotes the best performance obtained by a particular method for a particular classification task. Bold typeface denotes the best-performing method for the particular classification task.

We note that while VAEs lead to more accurate predictions, this performance improvement is not significant when compared to PCA. We conjecture that this might be an artefact of many linear relations present in the data, which are captured by the PCA. In contrast, the integrative VAEs are also able to model the non-linearities in the data, which gives them a performance advantage.

Comparing the performance of the four VAE architectures, H-VAE and X-VAE mostly yield more accurate predictions, however, the difference is not significant. Overall, for these three tasks, H-VAE produces more stable and better quality predictions when applied for integrating clinical and mRNA data, given the design decisions outlined previously. While for simplicity we made the same design choices for all architectures, the performance of these models can be further improved, with careful calibration of both the architecture components as well as the hyper-parameters of the classifier considered.

### 3.3 Qualitative analyses

In the last set of experiments, we visually inspected the learned representations of the whole data set, obtained from the H-VAE by integrating clinical/mRNA data. Using tSNE diagrams, shown in Figure 7, we compared the level of disentanglement of the embedded data with both, raw (uncompressed) data as well as PCA-transformed data. The tSNE projections clearly show that H-VAE is able to produce more sparse and disentangled representations in comparison to raw or PCA transformed data. Note that the t-SNE projections of the raw and PCA-transformed data also indicate data separability. This may explain the competitive performance produced by the benchmark classifiers in the previous section, as well as the advantage of integrating clinical and mRNA data.

**Figure 7:**
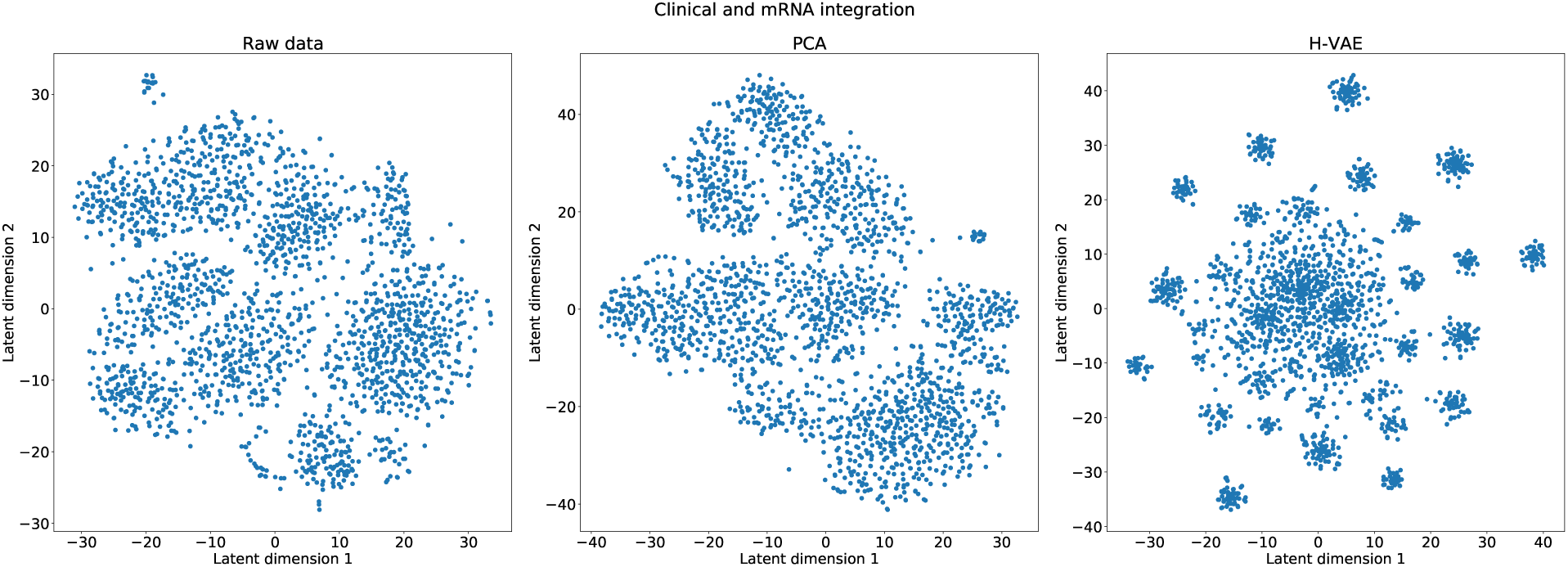
Qualitative comparison of the learned representations with H-VAE, raw data and PCA-transformed data when integrating clinical and mRNA data.

## Discussion

In this study we investigated and evaluated aspects of VAE architectures important for integrative data analyses. We designed and implemented four integrative VAE architectures, and demonstrated their utility in integrating multi-omics and clinical breast-cancer data. We systematically experimented (we evaluated 1296 different network configurations) with how the data should be integrated as well as what appropriate architecture parameters produce high-quality, low-dimensional representations. In the case of integrating breast-cancer data we found that the choice of an appropriate regularisation when training the autoencoders is imperative. Our results show that the integrative VAEs yield better (and more disentangled) representations when MMD is employed, which also corresponds to findings from other studies (Zhao et al., 2017; Chen et al., 2018). Moreover, we found that giving a moderately large weight to this regularisation term further improves the quality of the learned representations. The results show that the quality of the representations is mostly invariant to the size of the hidden layers and the embedding dimension, suggesting that the investigated integrative architectures are robust. Note however, that such parameters are task-specific, and therefore it is recommended that they are tuned according to the dimensionality of the input data as well as the depth of the network.

In the context of performance, all four integrative VAE architectures are generally able to produce better representations of the data when compared to a linear transformation approach. This suggests that the integrative VAEs are able to accurately model the non-linearities present in the integrated data, while still being able to reduce the data-dimensionality, leading to good representations. When comparing the different architectures, the results showed that overall the H-VAE and X-VAE exhibit the best performance, followed by the simple CNC-VAE and MM-VAE. This indicates that, while all of the architectures are able to accurately model the data, H-VAE exhibits more stable behaviour. Moreover, given that H-VAE is a hierarchical model, all of the learned representations (including the intermediate ones from the low-level autoencoders) can be further utilised for more delicate, interpretable analyses. Note however, when employing H-VAE, there is a trade-off between the quality of the learned representations and the time required for learning them. Therefore, when time or resources are limited, employing X-VAE or even the simple CNC-VAE will yield favourable results.

In terms of integrative analyses of breast-cancer data, the results indicate that, for the particular classification tasks considered in our study, some data types are more amenable to integrating than others. More specifically, utilising the VAEs for integrating clinical and mRNA data coupled with the right classification method led to better downstream predictive performance than the alternative integration of CNA and mRNA data. This highlights an important aspect of this study: for premium results in such integrative data analyses, one should not only focus on the choice and tuning of appropriate predictive methods, but also on the type of data at input. In other words, rather than considering separate components of the analysis, one should focus on the whole end-to-end integrative process.

Autoencoders have been used for learning representations and analysing transcriptomic cancer data before. In particular, our work relates to Way and Greene (2018), since it employs VAEs for constructing latent representations and analysing transcriptomic cancer data. The authors show that VAEs can be utilised for knowledge extraction from gene expression pan-cancer TCGA data (TCGA et al., 2013), thus reducing the dimensionality of the single, homogeneous data source while still being able to identify patterns related to different cancer types. Our work is also related to Tan et al. (2015), where the authors deploy Denoising Autoencoders (DAE) for integrating and analysing gene-expression data from TCGA (TCGA et al., 2013) and METABRIC (Curtis et al., 2012). Tan et al. (2015) also employ DAE for learning latent features from multiple data sets. The latent features are used to identify genes relevant to two different breast cancer sub-types.

In contrast to Curtis et al. (2012) and Tan et al. (2015), we designed novel VAE architectures for integrating heterogeneous data, hence enabling learning patterns that relate to the intrinsic relationships between different data types. While DAEs aim at learning an embedded representation of the input, the VAEs focus on learning the underlying distribution of the input data. Therefore, besides data integration, the methods proposed in this paper can be also employed for data generation.

More generally, our work relates to other approaches based on autoencoders for data integration on various tasks of cancer diagnosis and survival analysis. These include using Denoising Autoencoders for integrating various types of electronic health records (Miotto et al., 2016) as well as custom designed autoencoders for analyses of liver (Chaudhary et al., 2018), bladder (Poirion et al., 2018) and neuroblastoma (Zhang et al., 2018) cancer types.

In a broader context, our work is related to the long tradition of data integration approaches for addressing various challenges in cancer analyses. In particular, Curtis et al. (2012) present an approach for clustering breast-cancer patients based on integrated data from the METABRIC cohort. The approach uses the Integrative Clustering (IC) method (Shen et al., 2009) which produces clusters from a multi-omic joint latent embedding. These clusters are then utilised for identifying mutation-driver genes (Pereira et al., 2016) and survival analyses (Rueda et al., 2019). In this context, the work presented in this paper can be readily applied to similar tasks. In particular, the integrative VAEs can be used to learn common representations of the heterogeneous data at input, which can then be used for constructing clusters that address the aforementioned analysis tasks. In contrast to the IC method, the integrative VAEs can handle high-dimensional data sources, which provide better integration and therefore may further improve the overall performance.

In a similar context, the Similarity Network Fusion (SFN) method by Wang et al. (2014) successfully addresses intermediate heterogeneous data integration for identifying cancer sub-types for various kinds of cancers including glioblastoma, breast, kidney and lung carcinoma. SFN first constructs graphs from the individual data sources, which are in turn combined into a single, integrative, graph using nonlinear similarity approach. Such graphs can be also used in conjunction with the integrative VAEs. More specifically, by using such graphs will impose a structure of the integrative data, which in turn may lead to far better (and disentangled) representations. Next, Gevaert et al. (2006) present a data integration approach with Bayesian networks for predicting breast cancer prognosis. The authors report that employing Bayesian networks for intermediate integration yields better performance for the particular predictive task. Since our proposed VAE approaches address full data integration, they can also be readily used together with the aforementioned integrative approaches.

We identified several additional directions for future work. First, the experiments reported in this study are limited to integrating heterogeneous multi-omics data from two sources. While in principle the autoencoder designs allow for integrating heterogeneous data from many more sources simultaneously, we intend to empirically evaluate the generality of our approaches and extend them to other types of data such as imaging data. Next, considering the specific architecture decisions made in this paper, we plan to further refine the designed architecture and fine-tune the learning hyper-parameters in ordered to improve the quality of the learned representations. This includes experimenting with deeper architectures as well as implementing methods that allow for more sophisticated priors as well as methods that focus on more flexible posteriors (Rezende and Mohamed, 2015; Kingma et al., 2016). Finally, we intend to ensemble the various proposed architectures which should yield more stable and robust findings, and take a step further towards producing more meaningful and interpretable findings.

While VAEs are capable of generating useful representations for vast amounts of complex heterogeneous data, in terms of interpretability, the biological relevance of the learned representations has to be verified if they are to be used in clinical decision support systems. Previous work (Tan et al., 2015) has attempted to interpret latent features, wherein features which were most influential in deciding clinical phenomena such ER/IHC status were extracted and identified. However, the actual interpretations of these features have received comparatively little attention. In order to interpret extracted VAE features and bring explanation to the learned representations, biological and biomedical ontologies such as gene ontology (GO^3^) have proven very useful (Way and Greene, 2018; Titus et al., 2018). An immediate continuation of the work presented in this paper is performing enrichment analysis on genes most related to each VAEs’ learned embedding to investigate the joint effects of various gene sets within specific biological pathways. Tools such as ShinyGo^4^ allow KEGG Pathway Mapping^5^, where the relationships between genes and human disease including various types of cancer can be identified. Using this approach to interpretability can potentially offer a qualitative metric to evaluate and compare different VAE architectures based on the biological relevance of the features extracted from learned representations to breast cancer and other cancer types in general.

## Conclusion

In conclusion, in this study we demonstrate the utility of Variational Autoencoders for full data integration. The design and the analyses of different integrative VAE architectures and configurations, and in particular their application to the tasks of integrative modelling and analysing heterogeneous breast cancer data, are the main contributions of this paper.

The studied approaches have several distinguishing properties. First, they are able to produce representations that capture the structure (i.e., intrinsic relationships between the data variables) of the data and therefore allow for more accurate downstream analyses. Second, they are able to reduce the dimensionality of the input data without loss of quality or performance. Therefore, in the process of compressing the input data, they can reduce noise implicitly present in the data. Third, they are modular and easily extendable to handle integration of a multitude of heterogeneous data sets. Next, while the integrative VAEs can be used as a data pre-proccessing approach for learning representations, they can also be utilised in a more generative setting for producing surrogate data, which can be used for more in-depth analysis. Finally, we show that VAEs can be successfully applied to learn representations in complex integrative tasks, such as integrative analyses of breast cancer data, that ultimately lead to more accurate and stable diagnoses.

## Acknowledgments

This work was supported by The Mark Foundation for Cancer Research and Cancer Research UK Cambridge Centre [C9685/A25177]. We thank Dr. Jean Abraham and Dr. Oscar Rueda for the helpful feedback and discussions on the work presented in this paper.

## Supplementary

We investigated 108 different configurations for each of the four integrative architectures for each integrative task. Figure 8 and Figure 9 highlight the effect the optimal architecture designs when integrating CNA and mRNA data as well as clinical and CNA data, respectively. In particular, we investigate the size of the hidden dense layers, the size of the learned representation, the most appropriate regularisation in the objective function as well as how much this regularisation should influence the overall loss. These configurations are evaluated by comparing the average train and test performance of classifying IHC sub-typed patients.

**Figure 8:**
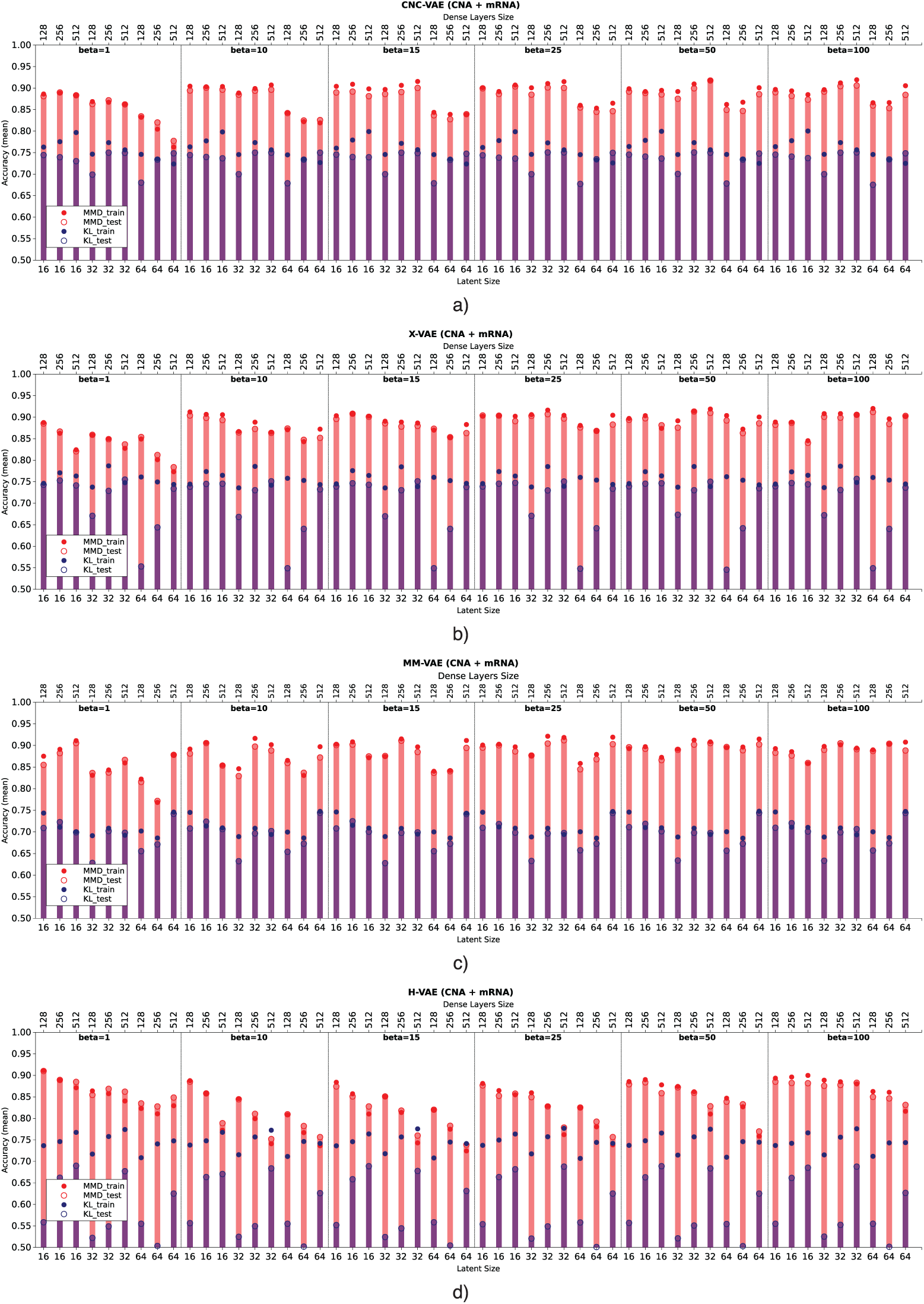
Comparison of the downstream performance on the IHC classification tasks of a predictive model trained on the representations produced by integrating CNA and mRNA data using (a) CNC-VAE (b) X-VAE (c) MM-VAE and (d) H-VAE. Full circles denote the training accuracy, while empty circles and bars denote the test accuracy averaged over 5-fold cross-validation. Red and blue colours denote the configurations when MMD and KL are employed, respectively. Bottom x-axis depicts the size of the latent dimension, while the top x-axis the size of the dense layers of each configuration.

**Figure 9:**
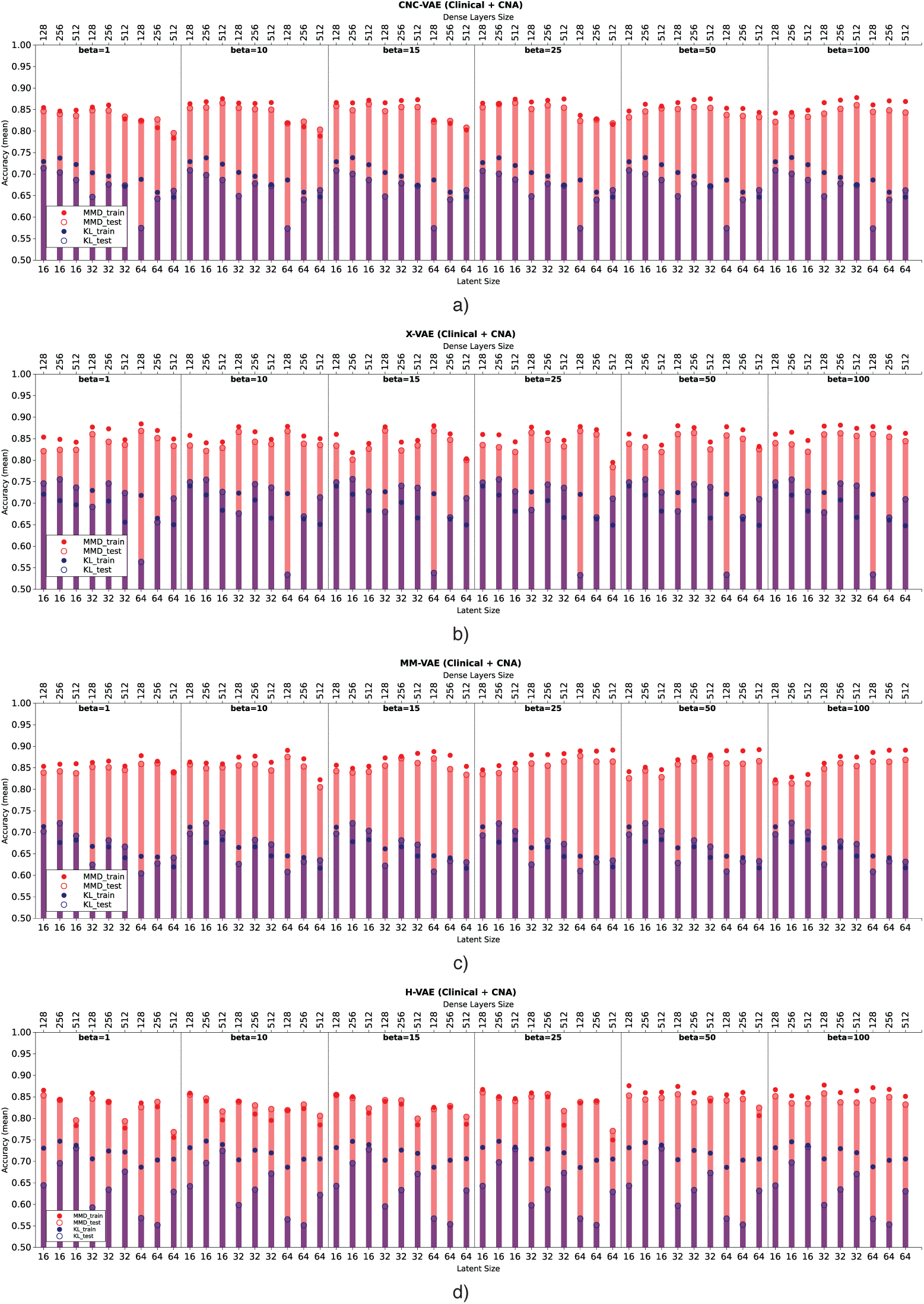
Comparison of the downstream performance on the IHC classification tasks of a predictive model trained on the representations produced by integrating clinical and CNA data using (a) CNC-VAE (b) X-VAE (c) MM-VAE and (d) H-VAE. Full circles denote the training accuracy, while empty circles and bars denote the test accuracy averaged over 5-fold cross-validation. Red and blue colours denote the configurations when MMD and KL are employed, respectively. Bottom x-axis depicts the size of the latent dimension, while the top x-axis the size of the dense layers of each configuration.

## Footnotes

2 Available after the completion of the review process. Please contact the authors.

3 http://geneontology.org

4 http://bioinformatics.sdstate.edu/go/

5 https://www.genome.jp/kegg/pathway.html#mapping

